# Tr1 cell-mediated protection against autoimmune disease by intranasal administration of a fusion protein targeting cDC1 cells

**DOI:** 10.1101/2023.04.11.536382

**Authors:** Charlotta Hansson, Cristina Lebrero-Fernández, Karin Schön, Davide Angeletti, Nils Lycke

## Abstract

Curative therapies against autoimmune diseases are lacking. Indeed, most of currently available treatments are only targeting symptoms. We have developed a novel strategy for a therapeutic vaccine against autoimmune diseases based on intranasal administration of a fusion protein tolerogen, which consists of a mutant, enzymatically inactive, cholera toxin A1-subunit genetically fused to disease relevant high affinity peptides and a dimer of D-fragments from protein A. The CTA1R7K-MOG/PLP-DD fusion proteins effectively reduced clinical symptoms in the experimental autoimmune encephalitis (EAE)-model of multiple sclerosis (MS). The treatment induced Tr1 cells, in the draining lymph node, which produced IL-10 and suppressed effector CD4^+^ T cell responses. This effect was dependent on IL-27 signalling, since treatment was ineffective in bone marrow chimeras lacking IL-27Rα within their hematopoietic compartment. scRNA-seq of dendritic cells (DC) in draining lymph nodes demonstrated distinct gene transcriptional changes of cDC1, including enhanced lipid metabolic pathways, induced by the tolerogenic fusion protein. Thus, our results with the tolerogenic fusion protein demonstrates the possibility to vaccinate and protect against disease progression by reinstating tolerance in MS and other autoimmune diseases.

## INTRODUCTION

There is a lack of curative therapies against autoimmune diseases ^1^. Nevertheless, many diverse treatments have been explored to dampen or potentially cure disease in experimental animal models, but few have been taken into the clinic ^2–7^. This is also true for the experimental autoimmune encephalitis (EAE) mouse model, which has been extensively studied to find a cure for multiple sclerosis (MS) ^8, 9^. Most of these therapies have been developed to target the inflammation in the brain and as such do not eliminate the cause of disease, but rather temporarily alleviate symptoms ^10^. However, by reinstating tolerance against disease-relevant proteins, such as myelin basic protein (MBP), myelin oligodendrocyte glycoprotein (MOG) or proteolipid protein (PLP), specific immunosuppression can potentially be achieved in MS patients ^11–13^. Although these peptides and proteins have been administered in various formulations and via different routes in the EAE model, we still lack vectors that reliably stimulate tolerance and can be used in humans ^14^. For example, mucosal administration of tolerogenic protein formulations have been found effective, but none have been developed passed clinical phase I/II-trials, albeit oral tolerance has been partly successful as, for example, with cholera toxin B subunit (CTB)-conjugated proteins given orally to patients with autoimmune uveitis (Bechet’s disease) ^15^. Hence, in awaiting the results of further clinical studies, we still have mostly symptomatic, or disease modyfing medication available today to treat MS and other autoimmune diseases ^16^.

It is well documented that effector CD4^+^ T cell subsets involved in disease progression in the EAE model, in particular Th1 and Th17 cells. This is not least illustrated by the fact that knocking out IFN-γ exacerbates EAE, while mice lacking IL-23p19, RORγt and IL-6 are resistant to EAE ^17–20^. Conversely, much effort has gone into advancing our understanding of how regulatory T cells (Tregs) can prevent EAE-disease progression ^21^. For example, transfer of nTregs before induction of either active or passive induction of EAE impairs disease onset and progression ^22, 23^ and CD25^+^FoxP3^+^ Tregs are known to accumulate in the central nervous system (CNS) during normal EAE development which correlates with disease recovery, though they are not able to suppress the peak of EAE disease ^14, 24^. In addition, Tr1 cells have been shown to protect against EAE in several studies, primarily by IL-10 production ^25^. For example, transfer of Tr1 cells induced by PD-L1 and anti-CD3 resulted in prevention of active EAE ^26^. Similarly, retinoic acid as adjuvant potently induced Tr1-mediated suppression of Th1 and Th17 ^27^. Furthermore, it is by now well established that IL-27 induces Tr1 cells and protects against EAE by acting on T cells directly ^28–33^ as well as on the antigen presenting dendritic cells (DCs) ^34, 35^. An important line of research for achieving Tregs and tolerance-induction is the focus on DCs, of which the classical DCs (cDCs) have been found critically important ^36^.

These findings and others exemplify the complex network within which several functional subpopulations of regulatory T cells cooperate to suppress auto-aggressive effector responses. We have developed a tolerance-inducing platform technology based on the enzymatically inactive mutant of cholera toxin A1 subunit, the CTA1R7K-X-DD fusion protein ^37^. In previous studies we have demonstrated that intranasal (i.n) treatment with this fusion protein carrying a disease-relevant peptide could significantly ameliorate or even prevent disease in two different autoimmune models, the collagen-induced arthritis (CIA) and experimental autoimmune myasthenia gravis (EAMG) models ^37–39^. Therapeutic effects correlated with increased IL-10 production, decreased effector responses and increased numbers of circulating FoxP3^+^ Tregs. Here, we explored if our tolerogenic vector could act as a therapeutic agent in the murine EAE model. We also aimed to further dissect the phenotypic changes that occurred in the T helper cells during the course of disease progression and treatment.

In the present study, we showed that the CTA1R7K-DD vector was effective in inducing tolerance to incorporated disease-relevant peptides, both when administered in parallel with disease induction as well as therapeutically. The effect was dependent on expansion of Tr1 cells in the mediastinal lymph nodes (mLN) and IL-27Rα signalling. Further, by scRNA-seq we elucidated the transcriptional reprogramming of tolerogenic cDC1 within the mLN.

In summary, our study further characterized the T cell mediated mechanisms that enforce tolerance induction subsequent to treatment with the CTA1R7K-DD vector. Our findings may represent a way forward in the search for effective therapeutic treatments against MS.

## RESULTS

### Intranasal treatment with the mutant CTA1R7K-MOG_35-55_-DD fusion protein protects against EAE disease

In earlier work we successfully developed a tolerogenic fusion protein based on a mutation in the CTA1 enzyme and a disease-specific peptide to prophylactically and therapeutically treat EAMG in mice ^38^. Along the same principles we now constructed a fusion protein, CTA1R7K-MOG_35-55_-DD, that we tested for its tolerogenic potential in the EAE model of MS. To this end we administered the tolerogen intranasally (i.n) on days 0, 2 and 4 (induction phase) or on days 10, 12 and 14 (disease development phase) or in both intervals. EAE-disease was provoked by intradermal injections at the base of the tail with MOG_35-55_ peptide in CFA on day 0 (**Fig. 1A**) ^40^. Disease development scoring revealed that the tolerogen was effective only when given in both intervals and we observed significantly milder EAE disease in these mice compared to the controls (**Fig. 1B-C**). Importantly, 50% of the treated animals did not develop disease, which is in agreement with our previous findings with this tolerance-inducing platform (**Fig.1D**) ^38^. Of note, i.n. treatment with an empty CTA1R7K-DD vector was completely ineffective, indicating that tolerization was peptide-dependent (**Fig.1C-D**). Moreover, an equimolar dose of MOG_35-55_ peptide given alone had no protective effect (**Fig.1C-D**). Mice given the CTA1R7K-MOG_35-55_-DD tolerogen exhibited nearly 90% reduction in infiltrating CD4^+^ T cells in the CNS and much fewer IFN-γ and IL-17 producing effector CD4^+^ T cells were detected as compared to levels found in untreated mice (**Fig. 1E**). In particular, the dominating IFNγ-producing CD4^+^ T cells, demonstrated a dramatic reduction, but also IL-17 producing cells were fewer in the CNS (**Fig. 1E**). Moreover, there was also a reduction in FoxP3^+^ Tregs and Foxp3^m^LAG3^+^CD49b^+^ Tr1 regulatory cells in the CNS of treated EAE mice, albeit this was less pronounced (**Fig. 1E**). None of the control groups had significantly reduced CNS-infiltrating effector CD4^+^ T cell levels (**Fig. 1E**). Thus, i.n treatment with the CTA1R7K-MOG_35-55_-DD tolerogen prevented or strongly ameliorated EAE-disease development concomitant with a substantially reduced number of infiltrating effector CD4^+^ T cells in the CNS.

**Figure 1.**
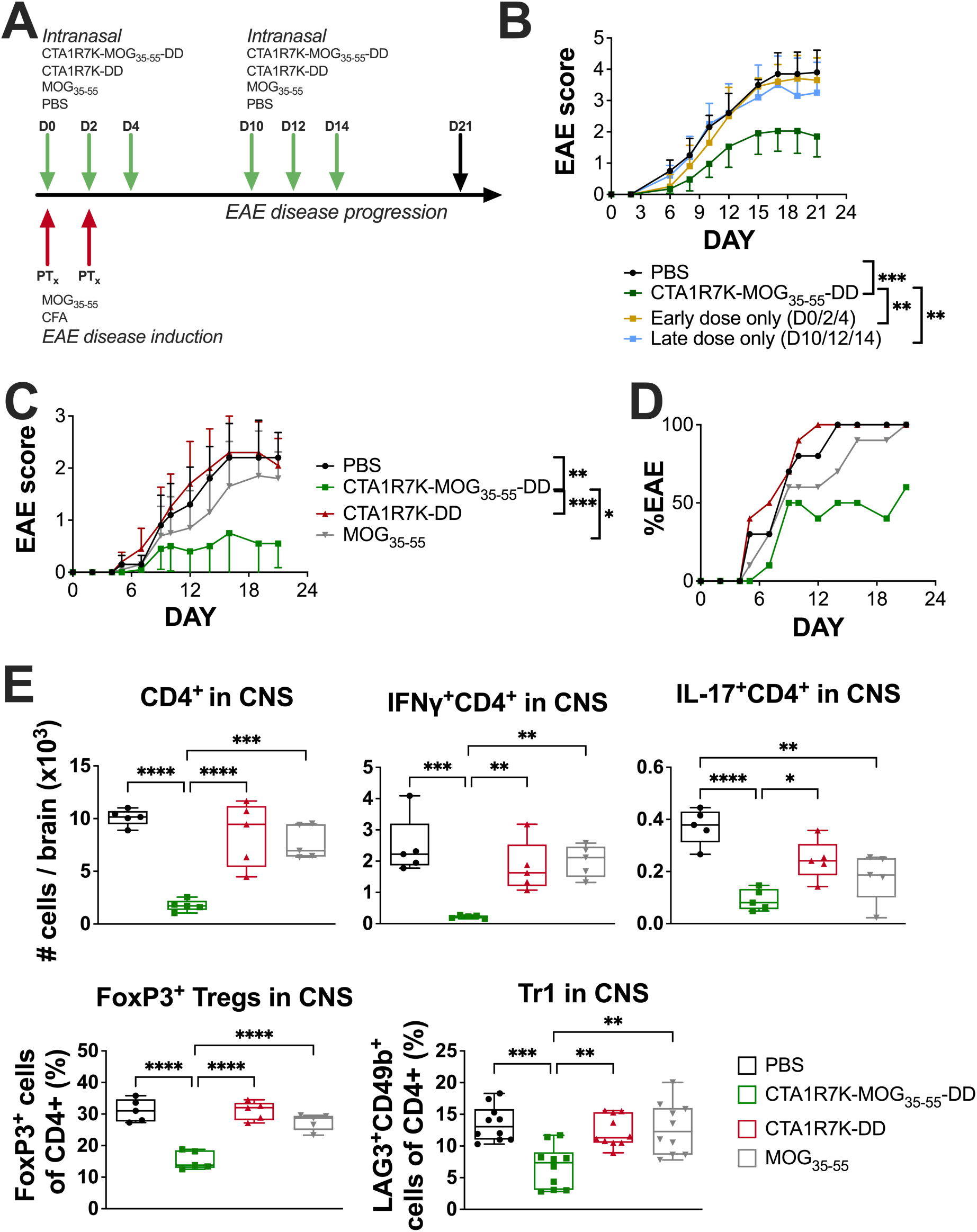
Intranasal treatment using the tolerogen CTA1R7K-MOG_35-55_-DD ameliorates EAE disease. **(a)** Schematic representation of EAE-disease induction and treatment (details in Materials & Methods section). EAE severity (**c**) and incidence (**d**) after treatment with PBS, MOG_35-55_ peptide, CTA1R7K-MOG_35-55_-DD or empty vector given at the normal dosing schedule (days 0, 2, 4, 10, 12, 14); or with CTA1R7K-MOG_35-55_-DD given only at early (induction phase: days 0, 2 and 4) or late (disease development phase: days 10, 12 and 14) time points (**d)** were assessed during the progression of the disease according to a 5-grade scale, and are shown as means±95% CI. Area under the curve (AUC) values were calculated for statistical analyses. **(e)** The absolute number of infiltrating CD4^+^ T cells per brain (meaning brain + spinal cord – central nervous system; CNS) producing IFN-γ or IL-17, and the frequency of FoxP3^+^ Tregs or Tr1 (CD49b^+^LAG3^+^) cells of all CD4^+^ T cells were determined by flow cytometry and values are given as means±SD. **(a, b, c, d, e)** The data is representative from at least two experiments (n=5-10). Statistical significance was determined by one-way ANOVA test followed by Tukey’s multiple comparison tests of every pair, and differences were considered statistically significant when P≤0.05 (*P≤0.05, **P ≤0.01, ***P≤0.001, and ****P≤0.0001).

### CTA1R7K-MOG_35-55_-DD effectively stimulates Tr1 cell development

Next, we evaluated if i.n treatment with the CTA1R7K-MOG_35-55_-DD tolerogen induced suppression of T cell responses in spleen (SP) and/or lymph nodes. Following treatment and EAE-induction we isolated CD4^+^ T cells from SP or inguinal lymph nodes (ILN) on day 21 and recall responses were assessed *in vitro* to a range of concentrations of MOG_35-55_ peptide. MOG-specific responses in both SP and ILN were substantially reduced, suggesting generalized immune suppression, in tolerized mice as compared to untreated control mice (**Fig. 2A**). Furthermore, all cytokines measured, IFN-γ, IL-17A, IL-17F, IL-22, IL-6, TNF-α and GM-CSF, were significantly suppressed (**Fig. 2B**). Hence, Th1 as well as Th17 effector CD4^+^ T cells in the lymph nodes and SP were effectively suppressed by i.n treatment with the tolerogen. A kinetic analysis demonstrated that already on day 11, representing the peak response in untreated mice, did we observe significantly reduced CNS-infiltrating CD4^+^ T cell numbers (**Fig. 2C**). Focusing on regulatory CD4^+^T cells we identified that Tr1 cells rather than Tregs (Foxp3^+^) were increased in the draining mediastinal lymph node (mLN) in tolerogen-treated mice, relative to what was observed in control mice (**Fig. 2D-E**). Thus, it appeared that the CTA1R7K-MOG_35-55_-DD tolerogen stimulated Tr1 cell development. When we sorted CD4^+^CD44^+^LAG3^+^CD49b^+^ Foxp3^m^Tr1 cells from mLN of treated mice these exerted strong suppression of MOG-peptide driven effector T cell proliferation, which strikingly also correlated with a 3-fold increase in IL-10 concentrations in these cultures (**Fig. 2F**). Taken together, i.n treatment with the tolerogen stimulated Tr1 cell development not only in draining lymph nodes but systemically in SP and ILN, which strongly suppressed MOG-specific effector CD4^+^ T cell responsiveness, greatly protecting against EAE disease development.

**Figure 2.**
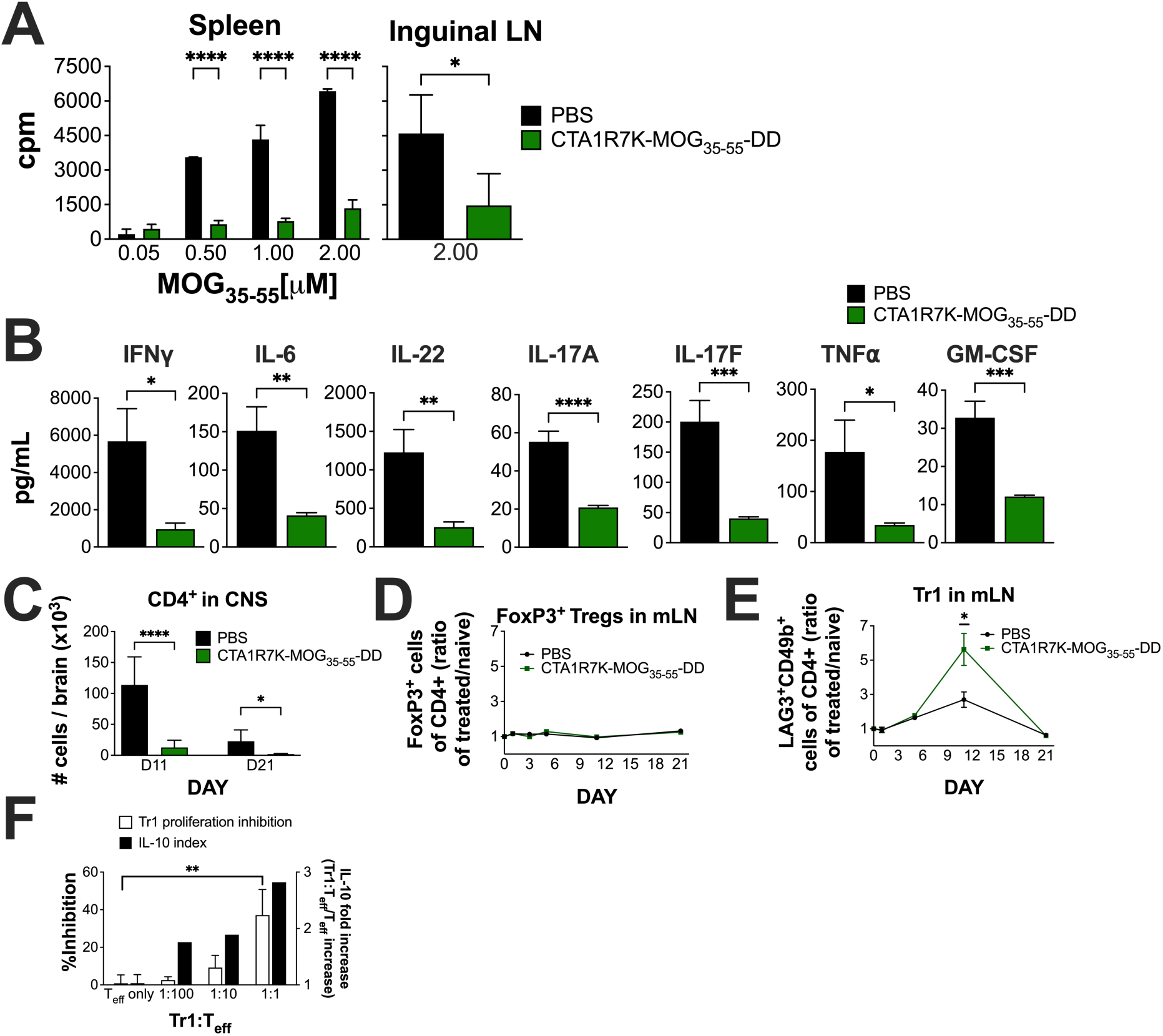
Intranasal tolerogen treatment greatly suppresses CD4^+^ T cell responses. **(a)** Isolated mononuclear cells from spleen and inguinal lymph nodes (ILN) were prepared on day 21 after induction of EAE disease. The recall response of CD4^+^ T cells to MOG_35-55_ peptide was determined *in vitro* with cells from mice treated i.n with CTA1R7K-MOG_35-55_-DD or untreated control mice. CD4^+^ T cell proliferation to a range of peptide doses in spleen or to a dose of 2μM of peptide in ILN was analysed. Values are given as mean cpms±SD of groups of 5 mice. **(b)** Indicated cytokines were assessed in supernatants from spleen and ILN CD4^+^ T cells from 5 mice per group stimulated with recall peptide at 2μM for 72h. Values are given for ILN in pg/ml as means±SD and are representative also of spleen cell cultures. **(a, b)** The data is representative of two experiments giving similar results. **(c)** The absolute number of infiltrating CD4^+^ T cells per brain (meaning brain + spinal cord – central nervous system; CNS) were determined at days 11 and 21 after EAE-disease induction; (**d**) the frequency (treatment/naïve ratio) of FoxP3^+^ Tregs or (**e**) Tr1 (CD49b^+^LAG3^+^) cells of all CD4^+^ T cells were quantified in mediastinal lymph nodes (mLN) over time. Data is pooled from at least two experiments (n=5-10) per time point and shown as means±SD. **(f)** For the inhibition assay, sorted Tr1 cells (CD4^+^CD44^+^LAG3^+^CD49b^+^) from the mLN of tolerogen-treated mice were co-cultured at different ratios with MOG-induced splenic effector T cells (CD4^+^CD44^+^LAG3^m^) from untreated mice, naïve CD11c^+^ APCs, and MOG_35-55_ peptide. Proliferation and IL-10 secretion in the culture supernatants were measured. Data represents one experiment with at least duplicates per condition and is shown as means±SD. Statistical significance was determined by unpaired two-tailed t-test, and differences were considered statistically significant when P≤0.05 (*P≤0.05, **P ≤0.01, ***P≤0.001, and ****P≤0.0001).

### MOG-specific TCR transgenic 2D2 cells exhibit Tr1 development and lack of effector 2D2 cells in CNS following i.n CTA1R7K-MOG_35-55_-DD treatment

Because the *in vitro* culture experiment clearly suggested that MOG-driven proliferation of effector CD4^+^ T cells in EAE mice was suppressed by Tr1 cells, we extended these studies to include TCR Tg MOG_35-55_-peptide specific 2D2 CD4^+^ T cells to get a more precise idea of the specificity of the Tr1 cells ^41^. To this end we adoptively transferred 2D2 cells into C57Bl/6 wild-type (WT) mice and monitored MOG_35-55_-specific CD4^+^ T cell migration into the CNS after EAE-induction. Tolerogen-treated mice had significantly reduced clinical scores, while mice receiving only 2D2-cells and untreated controls developed severe disease (**Fig. 3A**). Flow cytometric analysis of Tg Vα3.2^+^Vβ11^+^ TCR 2D2 CD4^+^ T cells from the CNS demonstrated that tolerized mice exhibited significantly fewer infiltrating CD4^+^ T cells in general and only few 2D2 T cells could be observed in i.n tolerogen-treated mice, of which a significant proportion were Tr1 cells (**Fig. 3B-C**), thus confirming that suppression was specific for pathogenic T cells.

**Figure 3.**
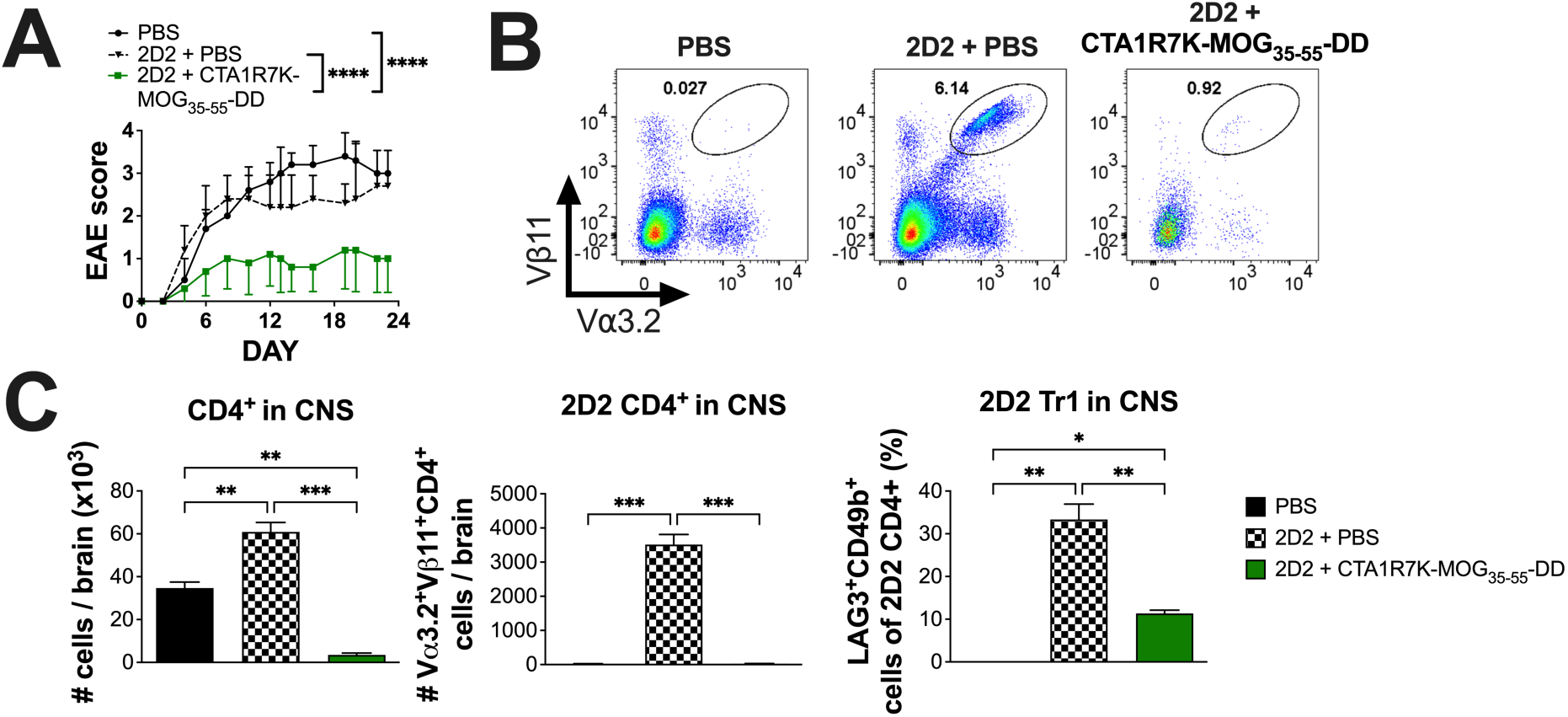
CTA1R7K-MOG_35-55_-DD treatment ameliorates EAE disease after 2D2 CD4^+^ T cells transfer *in vivo*. **(a)** 2D2 CD4^+^ T cells were freshly isolated and transferred to WT recipients prior to EAE-disease induction, and disease severity after treatment with PBS or CTA1R7K-MOG_35-55_-DD were assessed during the progression of the disease according to a 5-grade scale, and are shown as means±95% CI. Area under the curve (AUC) values were calculated for statistical analyses. **(b)** Central nervous system (brain + spinal cord; CNS) were analysed at the end of the experiment by flow cytometry and (**c**) absolute numbers of infiltrating CD4^+^ T cells and 2D2 (Vα3.2^+^Vβ11^+^) CD4^+^ T cells, and frequency of Tr1 (CD49b^+^LAG3^+^) cells of all 2D2 CD4^+^ T cells were quantified. Data is presented as means±SD and representative FACS plots are also shown. **(a, b, c)** Data is representative of two experiments (n=5). Statistical significance was determined by one-way ANOVA test followed by Tukey’s multiple comparison tests of every pair, and differences were considered statistically significant when P≤0.05 (*P≤0.05, **P ≤0.01, ***P≤0.001, and ****P≤0.0001).

### Therapeutic treatment with intranasal tolerogen blocks relapsing-remitting EAE-disease

To assess whether i.n tolerogen-treatment could be used therapeutically to prevent relapsing-remitting EAE disease, we used the well established SJL/J mouse model, which more closely resembles MS disease ^8^. Mice were treated i.n with CTA1R7K-PLP_139-151_-DD according to our established protocol and EAE was induced by PLP_139-151_ peptide injections in CFA. To test the therapeutic function we used a fusion protein with the PLP_178-191_ peptide, CTA1R7K-PLP_178-191_-DD, given i.n after the first peak of disease. Indeed we found that the PLP_139-151_-induced EAE SJL/J mouse model responded with greatly reduced disease development not only after prophylactic, but more importantly, to the therapeutic treatment with the CTA1R7K-PLP_178-191_-DD tolerogen, after the first peak of disease (**Fig. 4A**). Thus, despite significant EAE scores in the first peak, our tolerogen effectively blocked relapse of EAE disease in SJL/J mice (**Fig. 4A**). Interestingly, a cocktail combination PLP_139-151_ and PLP_178-191_ fusion proteins exerted a clear synergistic effect on preventing relapsing EAE disease, while PLP_139-151_ fusion protein alone had no therapeutic effect on relapsing disease (**Fig. 4B**). The use of an unrelated CTA1R7K-MOG_35-55_-DD fusion protein had no effect on PLP_139-151_-induced EAE-disease, indicating that cognate peptide-specificity is critical for the tolerizing effect (**Fig. 4C**). The presence of CNS-infiltrating effector CD4^+^ T cells in tolerogen-treated mice was monitored and we detected dramatically reduced cell numbers in tolerized mice (**Fig. 4D**). In fact, both IFN-γ and IL-17 producing effector CD4^+^ T cells were largely undetectable in the CNS (**Fig. 4d**).

**Figure 4.**
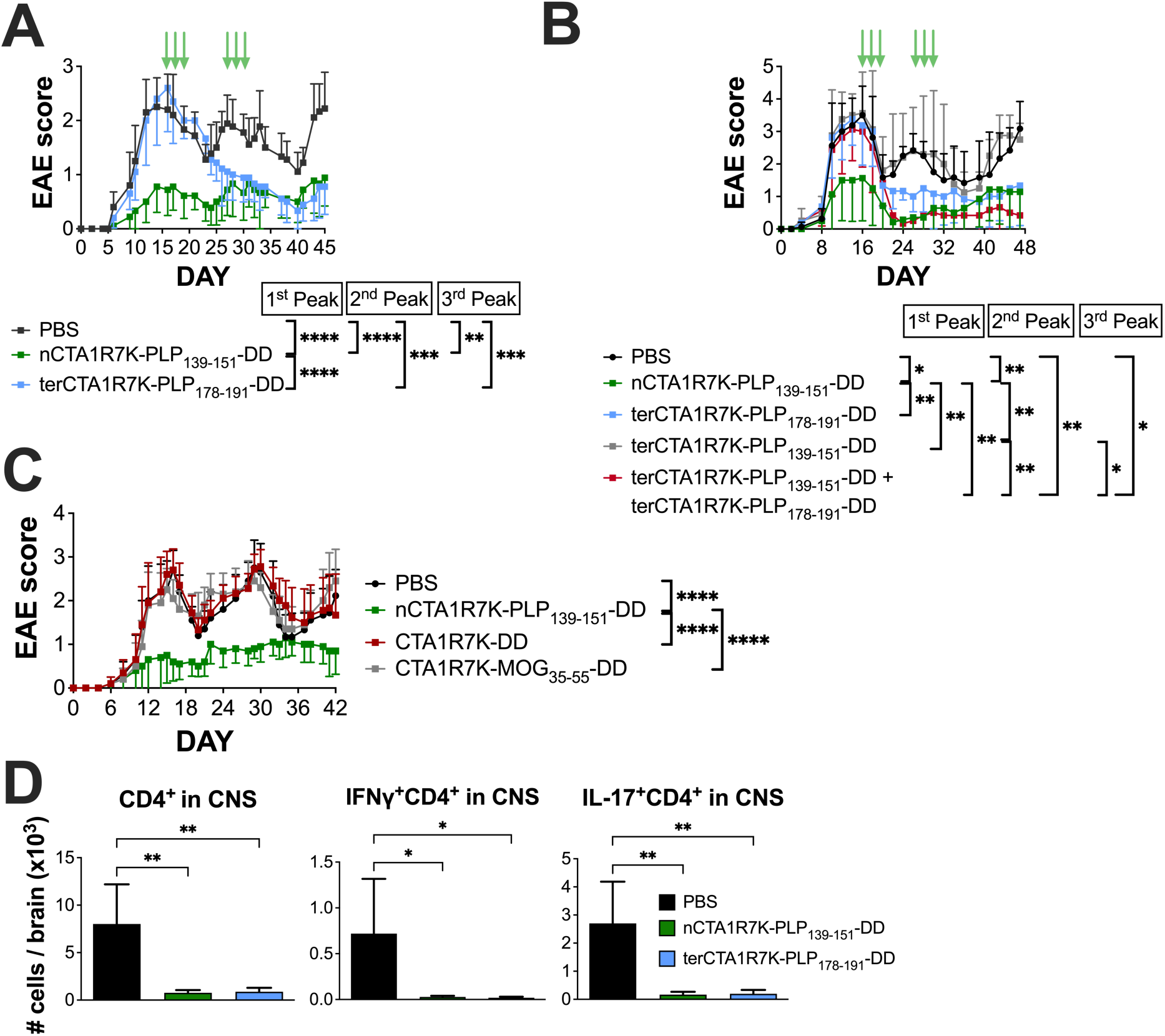
Therapeutic treatment is effective in a relapse-remitting EAE-model. EAE disease was induced in SJL/J mice using PLP_139-151_ peptide. **(a)** Mice were either treated as normal on days 0, 2, 4, 10, 12 and 14 (nCTA1R7K-PLP_139-151_-DD) or therapeutically shortly after the first disease peak (thCTA1R7K-PLP_178-191_-DD) as demonstrated by the green arrows in the graph. **(b)** Different PLP epitopes were tested for therapeutic treatment in PLP-induced EAE: terCTA1R7K-PLP_178-191_-DD, terCTA1R7K-PLP_139-151_-DD or a terCTA1R7K-PLP_139-151_-DD + terCTA1R7K-PLP_178-191_-DD cocktail, as compared to the normal nCTA1R7K-PLP_139-151_-DD dosing schedule. **(c)** Different constructs were used as treatment with the normal dosing schedule (days 0, 2, 4, 10, 12 and 14) in the PLP-induced EAE-model, namely CTA1R7K-PLP_139-151_-DD, CTA1R7K-DD and CTA1R7K-MOG_35-55_-DD. **(a-c)** Disease severity was assessed during the progression of the disease according to a 5-grade scale, and is shown as means±95% CI. Area under the curve (AUC) values were calculated for statistical analyses. Data is representative of at least two experiments (n=8-10). **(d)** At the end of the experiment presented in figure 4a, central nervous system (brain + spinal cord; CNS) were collected and analysed for infiltrating CD4^+^ T cells and IL-17 or IFN-1-expressing CD4^+^ T cells by flow cytometry. Data is shown as means±SD, and is representative of two experiments (n=5). Statistical significance was determined by one-way ANOVA test followed by Tukey’s multiple comparison tests of every pair, and differences were considered statistically significant when P≤0.05 (*P≤0.05, **P ≤0.01, ***P≤0.001, and ****P≤0.0001).

### Tolerogen-induced suppression by Tr1 cells is lost in IL-27Rα-/- mice following CTA1R7K-MOG_35-55_-DD treatment

Because IL-27R-signalling is critical for the generation of Tr1 cells we investigated to what extent mice lacking this receptor were able to respond to tolerization with the CTA1R7K-MOG_35-55_-DD fusion protein ^33, 34, 42–48^. We generated chimeric mice by transferring bone marrow from *wt* or *IL-27rα ^−/−^* mice into irradiated C57Bl/6 mice. This way we could focus specifically on CD4^+^ T cell differentiation requiring IL-27R-signalling. Following i.n treatment with CTA1R7K-MOG_35-55_-DD we failed to observe tolerance-induction in IL-27Rα-deficient chimeric mice (**Fig. 5A**). In contrast, WT bone marrow chimeric mice were effectively tolerized and exhibited significantly reduced EAE-disease levels (**Fig. 5A**). Of note, susceptibility to EAE disease in untreated mice was not significantly different between chimeric mice (**Fig. 5A**). Therefore, we concluded that IL-27R-signalling in CD4 T cells was required for tolerance-induction in EAE following i.n treatment with CTA1R7K-MOG_35-55_-DD, directly supporting the notion that Tr1-cells mediate suppression of effector CD4 T cells in our tolerogen-treated EAE model.

**Figure 5.**
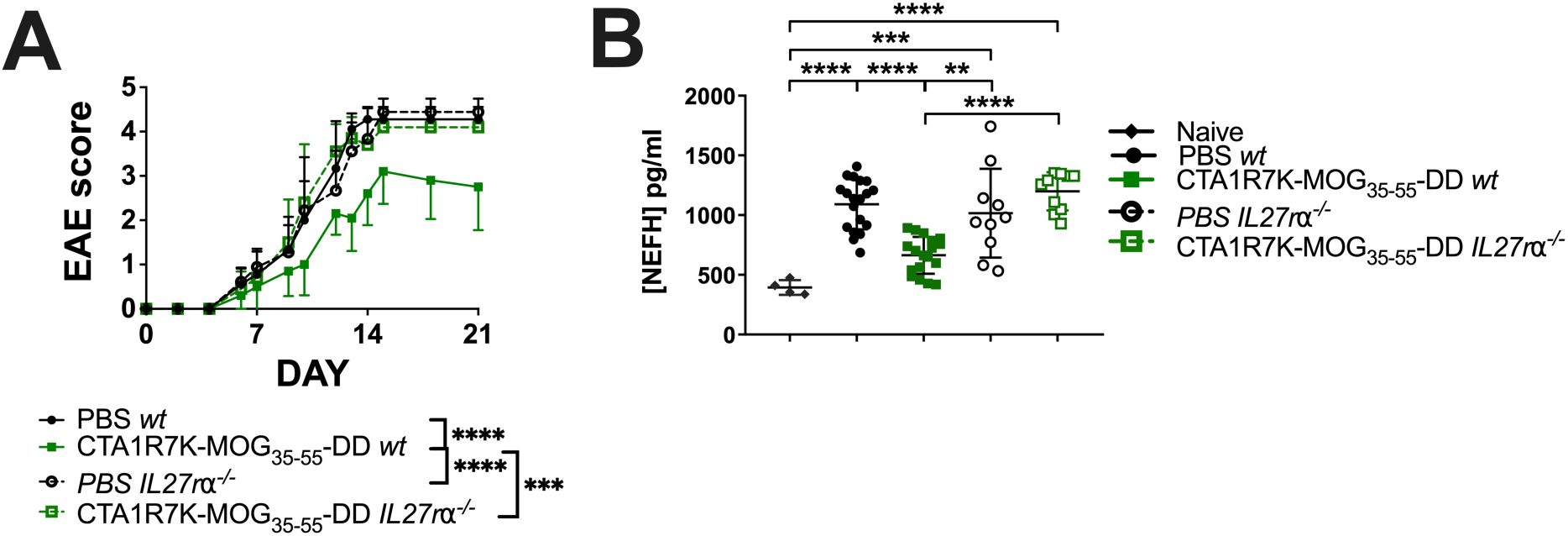
IL-27R-signalling is critical for EAE disease reduction after CTA1R7K-MOG_35-55_-DD treatment. EAE disease was induced in either [*wt* to *wt*] or [*IL-27rα^−/−^* to *wt*] bone marrow chimeras and treatment with CTA1R7K-MOG_35-55_-DD was carried out as previously indicated. **(a)** Disease severity was assessed during the progression of the disease according to a 5-grade scale, and is shown as means±95% CI. Area under the curve (AUC) values were calculated for statistical analyses. **(b)** Neurofilament Heavy Polypeptide (NEFH) levels were analysed in serum samples from naïve mice, and CTA1R7K-MOG_35-55_-DD-treated or untreated [*wt* to *wt*] or [*IL-27r-^−/−^* to *wt*] bone marrow chimeras by ELISA. Analysis of data and quantification was performed on the basis of corresponding standards curves. Data is shown as means±SD. Data is representative **(a)** or pooled **(b)** from at least two experiments (n=5-10). Statistical significance was determined by one-way ANOVA test followed by Tukey’s multiple comparison tests of every pair, and differences were considered statistically significant when P≤0.05 (*P≤0.05, **P ≤0.01, ***P≤0.001, and ****P≤0.0001).

These studies were extended to involve analysis of a recently established biomarker for MS disease, namely neurofilament light (NFL) in serum ^49, 50^. NFL levels have been strongly linked to the degree of MS/EAE-disease and nerve tissue damage ^51^. It was, therefore, striking to observe that tolerized WT mice had significantly lower serum NFL levels as opposed to bone marrow IL-27Rα-deficient chimeric mice following i.n treatment with CTA1R7K-MOG_35-55_-DD, the latter exhibiting levels comparable to untreated EAE control mice (**Fig. 5B**). Thus, mice deficient in the IL-27Rα CD4 T cells exhibited higher NFL serum levels and more dramatic tissue damage as they failed to respond to i.n treatment with our tolerogen, further supporting a critical role of Tr1 cells in CTA1R7K-MOG_35-55_-DD-induced tolerance (**Fig. 5B**).

### The CTA1R7K-MOG_35-55_-DD tolerogen binds to and acts through cDC1 cells which acquire a unique gene expression profile

In a previous study we demonstrated that fusion proteins with an active unmutated CTA1-enzyme exert strong immunoenhancing effects as they bind and are taken up by classical migratory DCs following i.n administration ^37, 52^. Interestingly, although the fusion proteins bind both cDC1 and cDC2 cells, we consistently found that only cDC1 cells were mediating the immunoenhancing effect following i.n administration, as also supported by the lack of an effect in Batf3^−/−^ mice, which do not have cDC1 cells ^52^. Therefore, to determine whether the mutant CTA1R7K-MOG_35-55_-DD acted via cDC1 cells we isolated cDCs from the mLN at 24h following i.n administration of fusion protein. To this end we used the CTA1R7K-DD and CTA1-DD fusion proteins, without incorporated peptide, to dissect if the tolerogen affected gene expression in targeted cDC cells differentlty to that of the CTA1-DD immunoenhancer ^52^. Dendritic cells from the mLN were FACS-sorted and subjected to single cell RNAseq (scRNAseq) analysis. We investigated to what extent migratory DCs changed gene expression profiles as a consequence of binding and up-take of of the tolerogen. The 10X Chromium platform technology was used and gene expression profiles were generated and compared with those of migratory DCs from unimmunized PBS-treated control mice ^53^. Of note, the isolated control cDCs from unimmunized mice represented naturally activated cDCs that have migrated to the mLN ^54^. After quality control and filtering we used the Seurat R toolkit to perform unsupervised clustering of the cDCs based on differentially expressed genes (DEG) and the data were visualized using uniform manifold approximation and projection (UMAP) in two dimensions (**Fig. 6A**) ^55^. We analyzed a minimum of 500 cells in each category. The migratory DCs were distributed in 5 clusters and a distinct separation of cDC1 and cDC2 cells into the clusters was evident using published gene lists defining the subsets (**Fig. 6B**) ^54^. Whereas migratory cDC2 cells clustered together irrespective of treatment the cDC1 cells clustered in distinct groups dominated by CTA1-, CTA1R7K- or PBS exposed cells (**Fig. 6C**). Using the Monocle 3 program we projected the cDC1 cells into a new UMAP and generated a pseudotime trajectory analysis that revealed striking differences between CTA1-DD and CTA1R7K-DD clusters, suggesting that the tolerogen had a unique and distinct impact on the targeted cDC1 cell, separate from that of the immunoenhancing enzymatically active CTA1-DD fusion protein (**Fig. 6D**). The two fusion proteins also clearly had distinctly different effects on cDCs than the naturally activated (PBS) cDC1 cells (**Fig. 6D**). Thus, cDC1 cells exposed to the tolerogen exhibited a gene expression profile that was unique and different; using Monocle 3 we analyzed signature genes, grouped into modules, by Louvain community analysis. This analysis demonstrated distinct modules representing cells that had been exposed to either the tolerogen, the immunoenhancer or the naturally activated cDC1 cells (**Fig. 6E**). The modules showing most dramatic changes were modules 2, 3, 4, 1 and 6 (**Fig. 6E**). In particular, genes in module 4 were highly expressed after exposure to the tolerogen compared to what was observed in CTA1-DD or PBS cDC1 cells (**Fig. 6E**). In this module we find genes linked to antigen presentation, cell signaling and lipid metabolism. Indeed, significantly upregulated genes characterizing tolerogen treatment of cDC1 cells were; *ccl17, Tm4sf5, cd74, Tnni2, Wim, S100a4, IL4i1, Syngr2, Tyrobp, Ifi30 and Myo1g* gene expression. We extended the analysis to include a gene ontology (GO) assessment of the upregulated pathways or processes for the tolerogen, the immunoenhancer or the naturally activated cDC1 cells (**Fig. 6F**).

**Figure 6.**
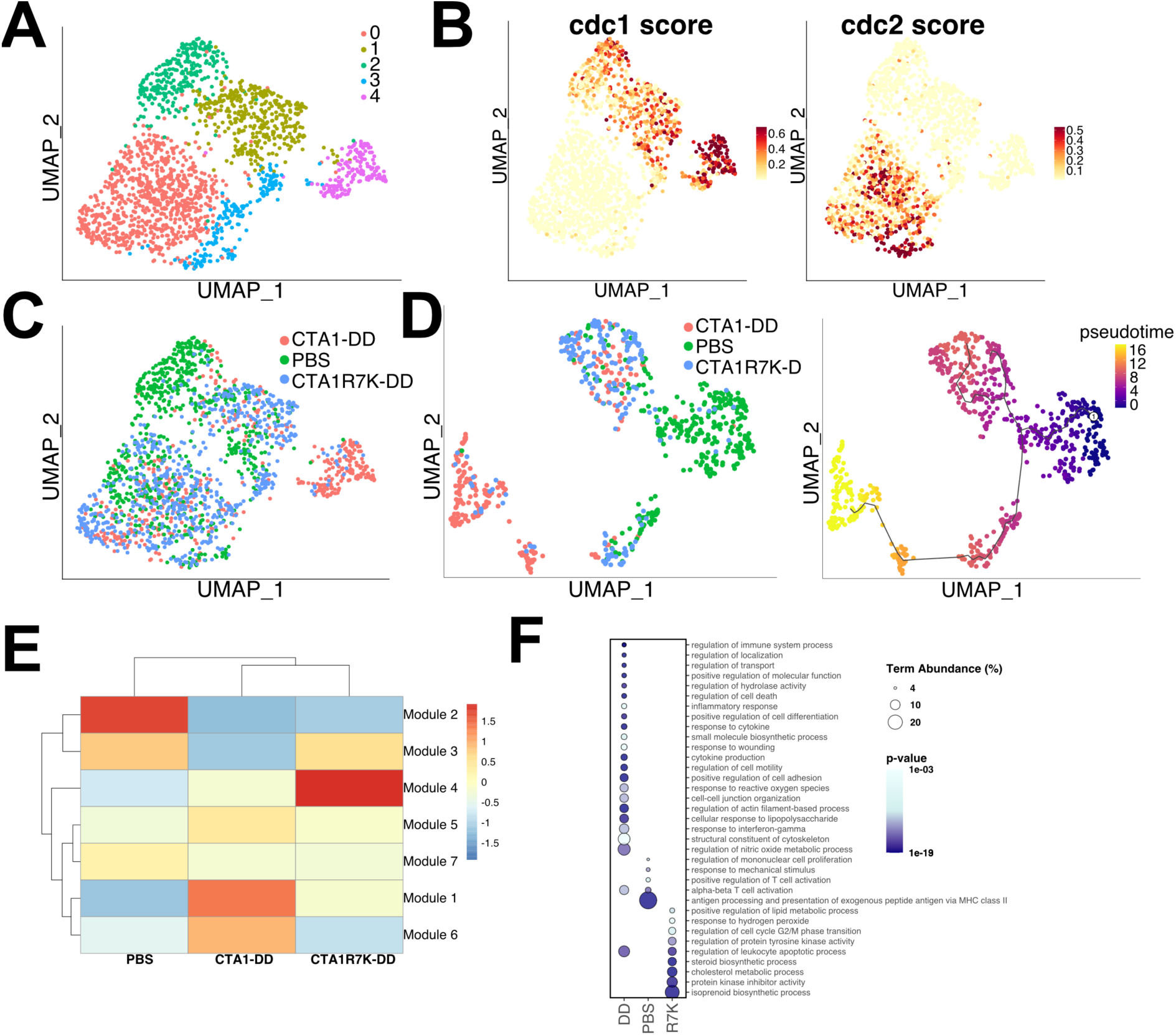
Single cell RNA sequence analysis of cDC1 cells subjected to the tolerance inducing CTA1R7K-MOG_35-55_-DD fusion protein. C57Bl/6 mice were immunized with CTA1R7K-MOG_35-55_-DD, CTA1-DD or PBS and migratory cDCs (MHC II ^high^ CDllc ^high^) sorted from mLN and subjected to scRNAseq. **a**) UMAP plot visualizing unsupervised clustering of all cells and treatment distribution of cells. **b**) Distribution of cDC1 and cDC2 cells based on expression of subset-specific gene sets. **c**) cDC1 cells were subsetted and reanalyzed using Monocle3 and visualized as UMAP projection. Pseudotime analysis, with starting node in the PBS group, was performed and data visualized on the UMAP. **d**) Heatmap representation of gene modules expression, grouped by treatment. **e**) Visualization of selected gene modules expression on the UMAP projection. **f**) Analysis of GO Biological and Immunological processes was performed using the ClueGO plugin in Cytoscape and the significant processes, for each treatment group, are visualized. Size of the dot indicates the % of term in the GO category, while color intensity represents the p value.

A remarkable observation was the effect of the tolerogen on lipid metabolism and cholesterol metabolism, in particular, but also protein kinase inhibition, and cell cycle regulation. As expected, the immunoenhancer had strong association with cell motility processes, response to cytokines and cell differentiation (**Fig. 6F**).

To conclude, the treatment with a cDC-targeted tolerogen had dramatic and unique effects on the transcriptome of cDC1 cells with distinctly different gene expression compared to that induced by the unmutated immunoenhancer, CTA1-DD. Whereas the tolerogen and the immunoenhancer exhibited distinct gene expression profiles in cDC1 cells these gene signatures also differed from those of naturally migrating (PBS) cDC1 cells. This result strongly argued for that the tolerogen had direct impact on gene transcription in the targeted cDC1 cells, involving several pathways and lipid metabolism, in particular, effects that were distinct and different from that of the enzymatically intact CTA1-DD immunoenhancer. Hence, the effect of mutant CTA1R7K-peptide-DD given i.n. could represent a novel mechanism of tolerance-induction, leading to differentiation of Tr1 cells and protection against autoimmune disease.

## DISCUSSION

In the present study we demonstrated that tolerance can effectively be induced by i.n immunization with the CTA1R7K-MOG/PLP-DD fusion proteins and protection against disease was observed in two models of murine EAE, reflecting an effect also during relapsing-remitting disease. Mice treated i.n with the fusion protein exhibited milder symptoms of disease and significantly reduced infiltrating Th17 and Th1 cells in the CNS, albeit this did not correlate with an increased proportion of regulatory T cells in CNS. Rather, the presence of regulatory T cells in the CNS was also reduced in the immunized and protected mice. Furthermore, CD4^+^ T cells in tolerized mice produced markedly lower levels of IFN-γ and IL-17 compared to unimmunized mice ^37^.

Because the generation of regulatory CD4^+^ T cells requires the expansion of activated naïve cells in the initial phase of the response in the draining lymph node, we were interested in characterizing the regulatory environment that promoted their differentiation. We identified IL-27R-signalling as a critical pathway in establishing the suppressive state that correlated with protection against disease. The latter is evidenced by the observation that tolerization was unsuccessful in *IL-27Rα^−/−^* chimeric mice, which strengthens the idea that Tr1 cells play a major role in the suppression of effector CD4^+^ T cell responses following immunizations with the CTA1R7K-X-DD fusion protein. We observed reduced CD4^+^ T cell responses in the draining lymph node, which coincided with significantly fewer infiltrating CD4^+^ T cells into the CNS in treated mice. Taken together, these observations suggested that the induced regulation most likely had occurred in the lymph node rather than in the CNS. Because disease is induced via priming of CD4^+^ T cells in the ILN and the site for inducing regulatory CD4^+^ T cells is the mLN, we followed the expansion of FoxP3^+^ Tregs as well as Tr1 cells at different time points in the ILN, SP, mLN, CNS and dCLN. We found a dramatic increase in Tr1 frequencies in mLN on day 11, during the progressive phase of disease. Odoardi *et al.* showed in rats that encephalitogenic CD4^+^ T cells pass through the lung parenchyma and recirculate through the mLN before reaching the effector site in the CNS ^56^. So, alternatively the effector CD4^+^ T cells pass through the mLN on their way to the CNS and their function could be suppressed in the mLN by the Tr1 cells. Moreover, our hypothesis raises the question whether the Tr1 cell expansion observed in the mLN arises from naïve CD4^+^ T cells or is a result of a phenotypic and functional shift in primed effector CD4^+^ T cells that impairs their ability to pass the blood brain barrier (BBB) and enter the CNS.

We did not observe an increase in FoxP3^+^ Tregs in the mLN that corresponded with increased numbers of Tr1 cells. However, because we gated on the whole CD4^+^TCRβ^+^ T cell population, there is a risk that small, but important, changes in antigen-specific Treg frequencies are occluded by the majority of non-specific CD4^+^ T cells. We have not yet ruled out the possibility that our treatment in the EAE model also induces FoxP3^+^ Tregs, similar to our observations from other models ^39^. There is a connection between Tr1 cells, IL-27 and FoxP3^+^ Tregs; however, it is not yet fully elucidated. Whereas some investigators report that IL-27 enhances the function of Tregs ^57^, others suggest that Tregs inhibit their own expansion by inducing IL-27 production in the APC which inhibits FoxP3 transcription via STAT3 signalling ^58^. In addition, IL-27 is implicated in Tr1 development ^44^. Nevertheless, it may well be that both FoxP3^+^ Tregs and Tr1 cells establish CTA1R7K-X-DD mediated tolerance in a cooperative fashion.

With our scRNA-seq analysis we focused on DC in mLN primed with either an immune-activating or tolerizing fusion protein. Importantly, we could mostly detect an effect in targeted cDC1 cells, as we previously reported ^52^. However, the transcriptional programme of cDC1 cells activated with the tolerogen *vs* the immunoenhancer *vs* naturally migrating cells was strikingly different. Pseudotime analysis suggested that the two induced transcriptional programs were distinct and mutually exclusive. Importantly, while the immunoenhancer had strong association with activation of immune processes, cell motility and other pro inflammatory processes, the tolerogen promoted protein kinase inhibitor activity and several processes linked with lipid metabolism. Protein kinase inhibitors have been linked to tolerogenic DCs in several studies ^59, 60^. More interesting are the lipid metabolic pathways, indeed metabolic states are a recognized factor which drive DC polarization ^61^. Cholesterol accumulation in DCs has been previously shown to be linked to a pro-inflammatory state ^62, 63^ however the cholesterol pathway also generates oxysterols, oxidative derivatives of cholesterol, which have in turn a tolerizing capacity. 7-ketocholesterols binds Ahr, which expression is induced by by IL-27, and in turn promotes development of Tr1 cells ^43, 64^. Further, the cholesterol efflux pathway has been identified as crucial to maintain tolerance in DC ^63^, therefore it is possible that increased lipid biosynthesis in tolerogenic DC is linked with secretion of oxysterols which can promote Tr1 polarization. It is plausible that presence of steroids in the milieu may promote a positive feedback loop and enhance the number of tolerogenic DC, as steroids are well known to induce these cells ^65, 66^. Our findings highlight remarkable differences in DC activation and polarization induced by a single amino acid mutation in the immunoenhancer CTA1. How this small change can have such dramatic effects needs to be elucidated in further studies.

We used the PLP-induced EAE model to examine whether our fusion protein was able to mediate protection against relapsing-remitting disease. The different disease progression is a result of epitope drift ^67^. Thus, at the later stages of EAE, disease peaks are dominated by epitopes different from the PLP_139-151_ peptide, which was incorporated in the first fusion protein we tested. While this fusion protein was effective at reducing EAE symptoms when used from the onset of disease, it failed to confer protection if used after the first peak of disease. By contrast, treatment with a vector containing the PLP_178-191_ peptide - which constitutes the immunodominant epitope during the second disease peak - was able to prevent all down-stream relapses. However, most importantly, a combination of the two was most effective. Inbred mouse strains do not recapitulate the HLA polymorphisms and heterogeneous peptide epitope repertoires within the human population, and thus is an important issue when translating animal data into a clinical setting. Further investigations on the use of treatment cocktails with combinations of peptide inserts in the CTA1R7K-X-DD platform, may be the way forward.

In summary, our results with the tolerogenic fusion protein convey optimism as to the possibility to protect against disease progression by reinstating tolerance in MS and other autoimmune diseases. Here, we provide evidence that our tolerogenic adjuvant, the CTA1R7K-X-DD fusion protein, mediates tolerance to incorporated peptides in an inflammatory context which depends on gene transcriptional changes in targeted cDC1 cells, IL-27 signalling and, possibly, Tr1 induction in the mLN.

## Acknowledgements

This paper is dedicated to the memory of Nils Lycke, a wonderful friend, colleague and inspiring mentor. The study received funding from the People Programme (Marie Curie Actions) of the European Union’s Seventh Framework Programme FP7/2007-2013/ under REA grant agreement number 607690 to N.L.. It was also supported in parts by research funds from the Knut and Alice Wallenberg Foundation KAW 2013.0030, the Swedish Foundation for Strategic Research SB12-0088, The Swedish Cancer Foundation, The Swedish Research Council, the EU project UNISEC, LUA/ALF ALFGBG-531021 and the Lundberg foundation (to N.L.). The authors would like to thank Jan-Olof Andersson and Richard Christison at MIVAC Development AB for production of the fusion proteins.

## Author contributions

CH, CLF and NL designed the study and planned the experiments. CH, CLF, AS and KS performed the experiments and analyzed the data. DA and CLF performed bioinformatics and modeling analysis. CH, CLF, DA and NL analyzed and interpreted the data. CH, CLF and DA made the figures. CH, CLF, DA and NL wrote the manuscript.

## MATERIALS & METHODS

### Antigens and adjuvants

Complete Freund’s adjuvant (CFA), MOG_35-55_, PLP_139-151_ and PLP_178-191_ were purchased from MD Biosciences (St Paul, USA). As previously described, CTA1R7K-DD, harbouring one copy of MOG_35-55_ (CTA1R7K-MOG_35-55_-DD), PLP_139-151_ (CTA1R7K-PLP_139-151_-DD) or PLP_178-191_ (CTA1R7K-PLP_178-191_-DD) were prepared, and lack of enzymatic activity was confirmed by an ADP-ribosylation assay, using CT and CTA1-DD as positive controls ^68, 69^.

### Mice and immunizations

All the experiments were conducted according to the protocols (Ethical permit numbers: 33/15 and 3066/20) approved by regional animal ethics committee in Gothenburg. They were housed in the specific pathogen free animal facility of Experimental Biomedicine Unit at the University of Gothenburg. C57BL/6- and SJL/J mice were purchased from Charles River, USA. 2D2 MOG-specific TCR transgenic mice and *IL-27rα^−/−^* mice (all on I-A^b^ C57BL/6 background) were kept and bred at the Laboratory for Experimental Biomedicine, Göteborg University (Göteborg Sweden). Mice were housed under specific pathogen-free conditions and were age- and sex-matched for all experiments. ***MOG_35-55_***: 5µg of CTA1R7K-MOG_35-55_-DD or CTA1R7K-DD, or equimolar amounts of MOG_35-55_ peptide was administered i.n on day 0, 2, 4, 10, 12 and 14 during EAE-disease progression. ***PLP_139-151_ and PLP_178-191_***: 5µg of CTA1R7K-PLP_139-151_-DD was administered i.n on day 0, 2, 4, 10, 12 and 14 during EAE-disease progression. For therapeutic treatments, different combinations of constructs, as indicated in the figure legends, were administered i.n at the first peak of disease.

*IL-27r^−/−^* bone marrow chimeric mice were generated in C57BL/6 mice after irradiation with 8 gy and reconstitution with 1×10^6^ cells bone marrow from either IL-27r^−/−^ or C57BL/6 mice WT control mice. The recipient mice were treated with antibiotics for two weeks and rested for six additional weeks before induction of EAE.

### EAE induction

EAE was induced in 8-9 weeks old C57BL/6 mice or SJL/J mice after an intradermal injection with 100µg myelin oligodendrocyte glycoprotein (MOG_35-55_) or proteolipid protein (PLP_139-151_) peptide (MD Biosciences) emulsified in Complete Freund’s Adjuvant (CFA; final concentration of 2mg/ml *Mycobacterium tuberculosis*; MD Biosciences). At day 0 and 2, 200ng of Pertussis Toxin (PT; List, USA) was administered intraperitoneally (i.p). Mice were examined for clinical symptoms regularly and scored as follows; 0: no disease; 1: limp tail; 2: limp tail and attenuated movement; 3: partial hind limb paralysis or severely affected movement; 4: complete hind limb paralysis; 5: moribund state or death. Mice with a score of 4 or higher during examination were euthanized. Studies were approved by the University of Gothenburg Ethics Committee for Animal Experimentation.

### Isolation of lymphocytes from the CNS

CNS infiltrates were isolated as previously described ^70^. Briefly, mice were perfused using PBS. The brain and spinal cord were removed, homogenized and loaded on a 30:37:70% Percoll gradient (GE Healthcare, Uppsala, Sweden). Interphase cells were collected and incubated at 37°C in 5% CO_2_ for 6 hours with 20ng/ml Phorbol 12-Myristate 13-Acetate (PMA, Sigma), 1µg/ml Ionomycon (Sigma) and 10µg Brefeldin A (Sigma) in Iscovés medium, before flow cytometry analysis.

### Cell culture, cell proliferation and cytokine assay

Single-cell suspensions from either spleen or lymph nodes were cultured in round-bottomed 96-well plates (Nunc, Roskilde, Denmark) in Iscove’s medium supplemented with 50μM 2-ME (Sigma), 1mM 1-glutamine (Biochrom), 50μg/ml gentamicin (Sigma) and 10% heat-inactivated FCS (Biochrom), for 72 hours at 37°C in 5% CO_2_. Proliferation was assessed by a beta-scintillation counter (Beckman Coulter, Turku, Finland) measuring titrated thymidine incorporation (PerkinElmer, Boston, USA). Culture supernatants were analysed for cytokine expression using the Th1/Th17 panel from Luminex technologies according to the manufacturer’s instructions. Plates were read in a Luminex MAGPIX instrument (Luminex Corporation). Analysis of data and quantification of cytokines was performed using the Luminex xPONENT Software (Luminex Corporation) on the basis of corresponding standards curves.

### Flow Cytometry

For all FACS analyses, cells were isolated and incubated for 5 minutes with the FcR blocking Ab (24G2) prior to staining. Live Dead Fixable Aqua Dead Cell Stain Kit (Invitrogen) was included to eliminate non-viable cells. The following antibodies (from Biolegend unless stated otherwise) were used for flow cytometry analyses: α-CD4:APC-H7 (GK1.5, BD Pharmigen), α-Vα3.2:FITC (RR3-16, eBioscience), α-Vβ11:perCP-eFluor710 (KT11), α-CD49:PE (HMα2), α-LAG3:biotin (C9B7W) and streptavidin:eFluor450 (eBioscience), α-CD44:BV570 (IM7), α-FoxP3:AlexaFluor647 (150D), α-IFN-γ:BV605 (XMG1.2), α-IL-17:FITC (eBio17B7, eBioscience) or α-TCRβ:FITC (H57597, BD Pharmigen). For intracellular staining, cells were fixed and permeabilized using the FoxP3 Staining Buffer Set (eBioscience). Intracellular IFN-γ and IL-17 expression was assessed after 6 hours of PMA/Ionomycin stimulation. Finally, cells were analysed using an LSR II flow cytometer or sorted using an LSR II Aria (BD Biosciences). FlowJo software was used for analysis (Tree Star).

### Adoptive cell transfer protocol

Spleen and lymph nodes from naïve 2D2 MOG-specific TCR transgenic mice were pooled and mashed through a nylon filter. CD4^+^ T cells were then enriched from the single-cell suspension using the “Big Easy” EasySep CD4^+^ T-cell negative isolation kit according to the manufacturer’s instructions (STEM-CELL, Grenoble, France). 1×10^6^ CD4^+^ T cells were subsequently transferred intravenously (i.v) into C57BL/6 recipients. EAE was induced 24 hours after transfer.

### ELISA

Neurofilament Heavy Polypeptide (NEFH) levels were analysed in mouse serum samples using the ELISA Kit for Neurofilament Heavy Polypeptide (Cloud-Clone Corp.) Plates were read in the 96-well microplate ELISA reader (Sunrise, Tecan). Analysis of data and quantification was performed on the basis of corresponding standards curves.

### Single cell RNAseq and Seurat and Monocle 3 analysis

Migratory cDCs (MHC II ^high^ CDllc ^high^) from naïve/PBS-treated or i.n fusion protein immunized mice were isolated from mLN from a pool of 7-8 C57Bl/6 mice and FACS-sorted on a Aria II cell sorter (BD Bioscience). Labeling of cells prior to sorting by flowcytometry (Fusion, BD) included the following specific anti-CD4, anti-CD8, anti-B220 and anti-F4/80, as described ^71^. Cells were sorted into IMDM, before being pelleted and resuspended in IMDM-10% FBS. Approximately 5000 sorted cells were loaded on the 10x Chromium Controller (10x Genomics). The scRNA-seq libraries were prepared following the user guide manual provided by the 10X Genomics company. Data from scRNA-seq of DC sorted samples were individually processed using **c**ell **r**anger **a**nalysis **p**ipeline (10X genomics platform) and reads aligned on the mm10 genome assembly. The output form Cell Ranger then further analysed with Seurat Seurat 3.0. Raw UMI count matrices were loaded and merged into a single Seurat object. Cells were discarded if they met any of the following; percentage of mitochondrial counts greater than 6 percent per cell, number of unique features either below 200 or above 3500, T cell genes (*‘Cd3d’, ‘Cd3e’, ‘Cd3g’, ‘Trac’, ‘Trbc1’, ‘Trbc2’*) greater than 0.1 percent and B cell genes (*‘Cd79a’, ‘Cd79b’, ‘Ms4a1’, ‘Cd19’*), greater than 0.1 percent. Gene counts were log normalized and mean centered and scaled by their standard deviation and the following variables regressed out : the number of UMI and the percentage of mitochondrial counts. Finally, cells were clustered and UMAP applied to further reduce dimensionality for visualization. For analysis using Monocle v3, the Seurat object was imported using the as.cell_data_set function from SeuratWrappers and further processed. Modules of co-regulated genes, differentially expressed between treatments, were identified using the function find_gene_modules using a list of Louvain resolution and selecting the one with highest value.

### GO analysis

Genes, belonging to unique modules (module 2 for PBS, module 4 for R7K and modules 1,5, and 6 for DD), were used for the analysis. GO analysis was performed using the ClueGO plugin of Cytoscape and with GO Biological and Immunological Processes as query ^72, 73^. We filtered based on GO terms with minimum 4 genes and 4% settings. Only pathways with *P* < 0.001 were considered. Common pathways were consolidated by clueGO. Pathways were exported and plotted using ggplot2 package.

### Statistical analysis

For clinical scores, the area under the curve (AUC) was calculated for each individual. Depending on the number of groups, significant differences were calculated using either a two-tailed unpaired student’s t-test or a one-way ANOVA test followed by Tukey’s multiple comparison tests using the Prism Software (GraphPad); p-values ≤0.05 were considered statistically significant.

